# Machine learning on subcortical brain features: A study of sample size efficiency for neurodegenerative disease classification

**DOI:** 10.64898/2026.09.15.751847

**Authors:** Yanghee Im, Melody J.Y. Kang, Boris A. Gutman, Sophia I. Thomopoulos, Paul M. Thompson, Christopher R.K. Ching, the Alzheimer’s Disease Neuroimaging Initiative

## Abstract

Subcortical brain alterations are a key feature of dementia disease progression. Machine learning (ML) has been applied widely to MRI-based brain features in dementia, where performance depends on the model choice, training data size, and input feature characteristics. Most studies compare ML models using a single training sample size. Here, we evaluate the sample-size efficiency of ML models based on subcortical gross volume and vertex-wise shape features for dementia stage classification using 2,511 samples in the Alzheimer’s Disease Neuroimaging Initiative (ADNI). Learning curves were generated for dementia vs. cognitively normal controls (CN), dementia vs. mild cognitive impairment (MCI), and MCI vs. CN across increasing training sample sizes. Classification performance improved when increasing sample size for all models, with late-fusion models consistently achieving the highest performance, and Logit-TVL1 outperforming the other shape-based models. Learning curve analysis showed that classification performance was driven by the training sample size and the magnitude of anatomical group differences, and can be used to optimize future model selection tasks in dementia and other brain disorders.

## 1 Introduction

Subcortical brain alterations are an early sign of Alzheimer’s disease (AD) and related dementias [1, 2]. Identification of individuals at risk of progression from healthy aging to MCI and dementia can help to facilitate accurate diagnosis and treatment plans. While gross volumetric brain measurements are commonly used [3–6], more advanced surface-based and three-dimensional (3D) shape features provide higher spatial resolution and can detect localized morphometric changes to functionally distinct subregions of structures not captured by traditional brain volume estimates [7–12]. Through finer mapping, shape analysis may better capture the link between pathophysiological brain changes and different illness phases, as in healthy aging, cognitive impairment, and dementia [13].

Machine learning (ML) methods have been trained on brain shape features for a variety of tasks, including diagnostic classification [14, 15]. Linear models are widely used as they are computationally efficient, relatively interpretable, and less prone to overfitting than more complex approaches when sample sizes are limited. The high dimensionality of vertex-wise shape features relative to available sample sizes requires the use of dimensionality reduction and regularization techniques. Several approaches have been proposed to improve performance in high-dimensional settings. Principal components analysis followed by linear discriminant analysis (PCA-LDA) reduces dimensionality before classification, Tikhonov-regularized linear discriminant analysis (LDA-Tikhonov) stabilizes covariance estimation through regularization, and logistic regression with Total Variation and L1 regularization (Logit-TVL1) incorporates sparsity and spatial smoothness into the model [16, 17].

When determining the best ML model for a task, performance can depend on the available training data size and the magnitude of anatomical group differences. Most neuroimaging studies evaluate ML model performance using a single training sample size and report the final classification accuracy. This makes it difficult to determine model performance across different training sample sizes, especially as more complex models may eventually outperform simpler ones, given enough training data [18]. As the scale of neuroimaging samples continues to grow through large-scale collaborative initiatives, such as the ENIGMA Consortium [19], ADNI [20], UK Biobank [21], understanding how ML performance changes with sample size can guide model selection and help estimate the amount of data required to achieve reliable predictions.

Recent work showed that learning curves, which describe how model performance changes with increasing training data, follows predictable scaling behavior [18]. Under this scaling-law framework, the relative performance of ML models is expected to depend on the model, the sample size, and the characteristics of the underlying signal.

Evaluating learning curves across different training sample sizes will improve ML model selection.

In this study, we test how training sample size influences ML classification of neurodegenerative disease diagnosis using MRI-derived subcortical brain volume and shape features. We tested three shape-based ML approaches (PCA-LDA, LDA-Tikhonov, and Logit-TVL1) and late-fusion models, and compared them with volume-based models. Classification performance was tested for three diagnostic comparisons: dementia versus cognitively normal controls (CN), dementia versus mild cognitive impairment (MCI), and MCI versus dementia. We first quantified anatomical group differences using vertex-wise linear regression (Cohen’s *d* effect size maps) and then evaluated model performance across a range of training sample sizes. Learning curves were constructed by varying the training sample size to test how predictive performance scaled with larger sample sizes. We hypothesized that: (1) classification performance would improve with increasing sample size for all models, (2) the rate of improvement would differ across diagnostic comparisons according to the magnitude of the group difference, (3) model rankings would vary as a function of sample size, providing an empirical test of scaling-law predictions for volume-based, shape-based, and fusion models, (4) fusion model will achieve the best performance as gross volumetric measures provide a low-variance summary of global atrophy that complements high-dimensional shape features and guides the regularized model.

## 2 Methods

### 2.1 Data and Preprocessing

#### Participant Demographics

3D T1-weighted (T1w) volumetric brain MRI scans from 417 individuals with dementia, 1156 with mild cognitive impairment (MCI) and 932 cognitive normal (CN) participants collected from 63 Alzheimer’s Disease Neuroimaging Initiative (ADNI; adni.loni.usc.edu) sites were included in the study. Participant demographics are summarized in **Table 1**.

**Table 1.** Demographic characteristics of participants in ADNI.

|  | CN<br>(N = 932) | MCI<br>(N = 1156) | Dementia<br>(N = 417) |
| --- | --- | --- | --- |
| Age Mean (SD) | 72.1 (6.7) | 72.7 (7.6) | 74.8 (7.8) |
| Sex (M/F) | 399/533 | 666/490 | 233/184 |
| MMSE score<br>Mean (SD) | 29.0 (1.1) | 27.6 (1.8) | 23.1 (2.1) |
| CDR-SB score<br>Mean (SD) | 0.04 (0.14) | 1.51 (0.92) | 4.37 (1.87) |

#### Gross Subcortical Volume Processing

Gross bilateral volumes were extracted from seven regions (hippocampus, amygdala, caudate, putamen, pallidum, thalamus, and nucleus accumbens) using FreeSurfer (v7.1.1).

#### Subcortical Shape Processing

The ENIGMA Subcortical Shape Pipeline [22–24] uses FreeSurfer subcortical volumes to derive a shape feature from seven bilateral subcortical shape models. These standardized shape features have been used in the largest analyses across a range of brain disorders from the ENIGMA Consortium [8– 12]. Briefly, the geometry of each subcortical structure is represented by a 3D triangular surface mesh, which is registered to the ENIGMA shape template via the Medial Demons Algorithm [25]. The alignment is optimized through medial model fitting as well as intrinsic shape characteristics. Once correspondence is established, a local thickness shape feature is calculated at each vertex:

#### Radial distance (RD)

The local “thickness” of a structure is characterized by RD, a vertex-wise measure derived from the minimum Euclidean distance between the surface and its medial curve *c*(*t*). For each surface vertex *P*∈*M*, the radial distance is defined as:

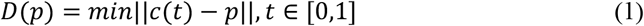

### 2.2 Group Difference Effect Size Estimates

#### Linear Mixed-Effect Model

To quantify gross volume and shape differences between diagnostic groups, and provide a statistical reference for interpreting ML performance and learning-curve behavior across diagnostic tasks, linear mixed model regression (LME) models were used adjusting for age, sex, intracranial volume (ICV), and scan site. Partial cohen’s *d* effect sizes were computed for each vertex-wise feature from the *t*-statistic for the diagnosis contrast [26]. Shape features were adjusted for covariates using a linear mixed effect model:

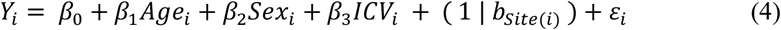

Where *Y*_*i*_ denotes for participant i, Age represents age at scan, Sex denotes biological sex, *ICV*_*i*_ indicates intracranial volume, (1|*b*_site(*i*)_) models random effect from site difference, and ε_*i*_ is the residual error term.

#### Linear Regression with ComBat-GAM

To adjust for potential site confounds (e.g., scanner field strength, acquisition protocol, etc.) we applied vertex-wise ComBat-GAM harmonization, a widely used extension of ComBat preserving non-linear age effects. We applied ComBat-GAM to the RD shape features using our Brain Geometry Toolkit [23, 24]. ComBat-GAM corrected shape features were also analyzed to investigate Cohen’s *d* effect sizes across data standardization techniques while adjusting for covariates using a linear regression model.

### 2.3 Machine Learning Models

#### Volume-Based Baseline Model

As a baseline for comparison with shape-based ML models, bilateral gross subcortical volumes were used as input features for a linear discriminant analysis (LDA) model.

#### Shape-Based Machine Learning Models

To evaluate the relationship between sample size and predictive performance, we implemented three linear shape-based ML models that differ in feature representation and regularization: PCA-LDA, Tikhonov-regularized LDA (LDA-Tikhonov), and logistic regression with Total Variation and L1 regularization (Logit-TVL1).

#### PCA-LDA

PCA-LDA reduces the dimensionality of the vertex-wise shape features prior to classification. PCA was applied to the training data to obtain a low-dimensional orthogonal representation of the original shape features. Principal components were estimated exclusively within each training fold, and the retained components explained a predefined proportion of the total variance. The reduced feature representation was then classified using LDA, which identifies a linear projection that maximizes between-class variance relative to within-class variance. By restricting the model to the dominant principal components, PCA-LDA emphasizes low-rank structure while reducing the influence of noise in high-dimensional feature spaces.

#### LDA-Tikhonov

Unlike PCA-LDA, the LDA-Tikhonov method operates directly on the original vertex-wise shape features. Classical LDA requires inversion of the within-class covariance matrix, which can become unstable when the number of features approaches or exceeds the number of training samples. Tikhonov regularization improves numerical stability by shrinking the covariance matrix toward the identity matrix,

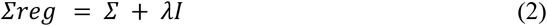

where Σ is the empirical covariance matrix, I is the identity matrix, and λ is the regularization parameter. This regularization reduces the influence of unstable covariance estimates while preserving the linear discriminant framework.

#### Logit-TVL1

Logistic regression with combined Total Variation (TV) and L1 regularization (Logit-TVL1) was used to incorporate sparsity and spatial regularization into the model. Model parameters were estimated by minimizing the logistic loss function with combined L1 and TV penalties. The optimization objective can be expressed as:

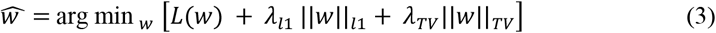

where L is the logistic loss, ||*w*||_*l*1_is the L1 penalty promoting sparse solutions, and ||*w*||_*TV*_ is the total variation penalty encouraging spatially contiguous coefficient patterns.

The L1 penalty promotes sparse solutions, whereas the TV penalty encourages neighboring vertices to have similar weights. This formulation allows the model to identify distributed shape patterns while maintaining interpretability. Compared to PCA-LDA and LDA-Tikhonov, Logit-TVL1 explicitly incorporates spatial information into the regularization of the model while retaining a linear decision boundary.

#### Shape-Volume Fusion Models

To assess whether conventional volumetric measures improve the sample-size efficiency of shape-based models, we also tested late fusion shape-volume models. Shape-based and volume-based models were trained independently, and their predictions were combined using a logistic regression meta-classifier.

### 2.4 Diagnostic Classification Tasks

Learning curves were computed separately for the three pairwise diagnostic classification tasks representing different stages along the healthy aging to dementia continuum: (1) dementia vs. CN, (2) dementia vs. MCI, and (3) MCI vs. CN. These comparisons were selected to reflect different levels of disease severity and expected classification difficulty.

Dementia vs. CN classification represents the largest expected effect size and therefore serves as a relatively high-signal classification problem. In contrast, MCI vs. CN and dementia vs. MCI involve more subtle morphological differences between groups, and are expected to require larger sample sizes to achieve comparable ML performance. Learning curves were estimated independently for each diagnostic comparison to test whether scaling behavior differed as a function of disease stage and classification difficulty.

### 2.5 Learning Curve Analysis

To test the relationship between sample size and classification performance, we performed a systematic learning curve analysis using subsampling ratios of 6.5%, 12.5%, 25%, 50%, 75%, and 100% of the available training data (70% of total sample), while leaving out 30% of total sample for independent test samples. Training and test data were divided randomly while preserving class proportions, and this procedure was repeated with 5 random draws. Learning curves were generated for PCA-LDA, LDA-Tikhonov, and Logit-TVL1 to evaluate how predictive performance scaled with increasing amounts of training data.

#### Subsampling Procedure

Bootstrapped resampling with 5 repeated random draws (iteration=500) was used at each sample size to reduce variability from a single random sampling and obtain robust estimates of model performance. For each bootstrap iteration, models were trained on the sampled training data and evaluated on the independently held out test data. Performance metrics were averaged across all bootstrap iterations and random draws. Variability was quantified using the distributions obtained from the bootstrap samples.

#### Cross-Validation

Model performance was evaluated using 3-fold stratified cross-validation within each subsampled dataset. Dimensionality reduction and hyperparameter optimization were performed within the training folds. Test data remained completely independent throughout model training and parameter selection.

#### Evaluation Metrics

We used the area under the receiver operating characteristic curve (AUC) as a threshold-independent measure of classification performance for the study. For each sample size, the mean and standard error of each metric were calculated across all bootstrapped samples.

#### Statistical Analysis

We compared AUC metrics of shape and fusion models to volume models using the paired bootstrap samples at each subsampling fraction for each task using a t-test. Statistical significance was established using false discovery rate (FDR, *q*<0.05) to correct for multiple comparisons.

## 3 Results

### 3.1 Gross Volume Group Differences

Those with dementia showed a pattern of smaller volumes across all seven subcortical regions except for lateral ventricles, which showed larger volumes compared to CN. This pattern was observed in the other group comparisons (dementia vs. MCI and MCI vs. CN), though caudate and globus pallidus were not significantly different after FDR correction for multiple comparisons.

### 3.2 Subcortical Shape Group Differences

Those with dementia showed a widespread pattern of lower thickness across most structures compared to MCI and CN (**Fig. 2**). While hippocampus, amygdala, and accumbens all showed primarily thinner patterns in dementia and MCI, the thalamus, caudate, putamen, and pallidum showed more complex shape alterations, including patches of locally greater (thicker) radial distance in those MCI or dementia compared to CN. LME models adjusting for site as a random effect showed a pattern of larger maximum effect sizes compared to models run on ComBat-GAM adjusted shape features.

**Fig. 1.**
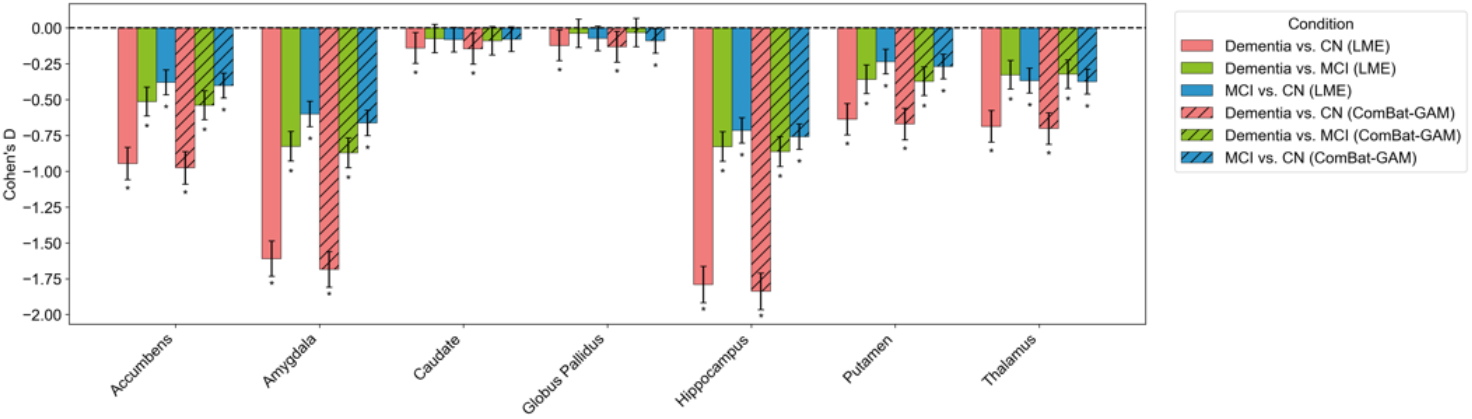
Cohen’s *d* effect size estimates from gross subcortical volumetric analysis using linear mixed-effects models (LME) adjusting for age, sex, intracranial volume (ICV), and scan site, and regression models based on ComBat-GAM adjusted gross volume features. All models were adjusted for age, sex and ICV. All significant group differences (*asterisk) were adjusted for multiple comparisons using the false discovery rate (FDR) (*q*<0.05).

**Fig. 2.**
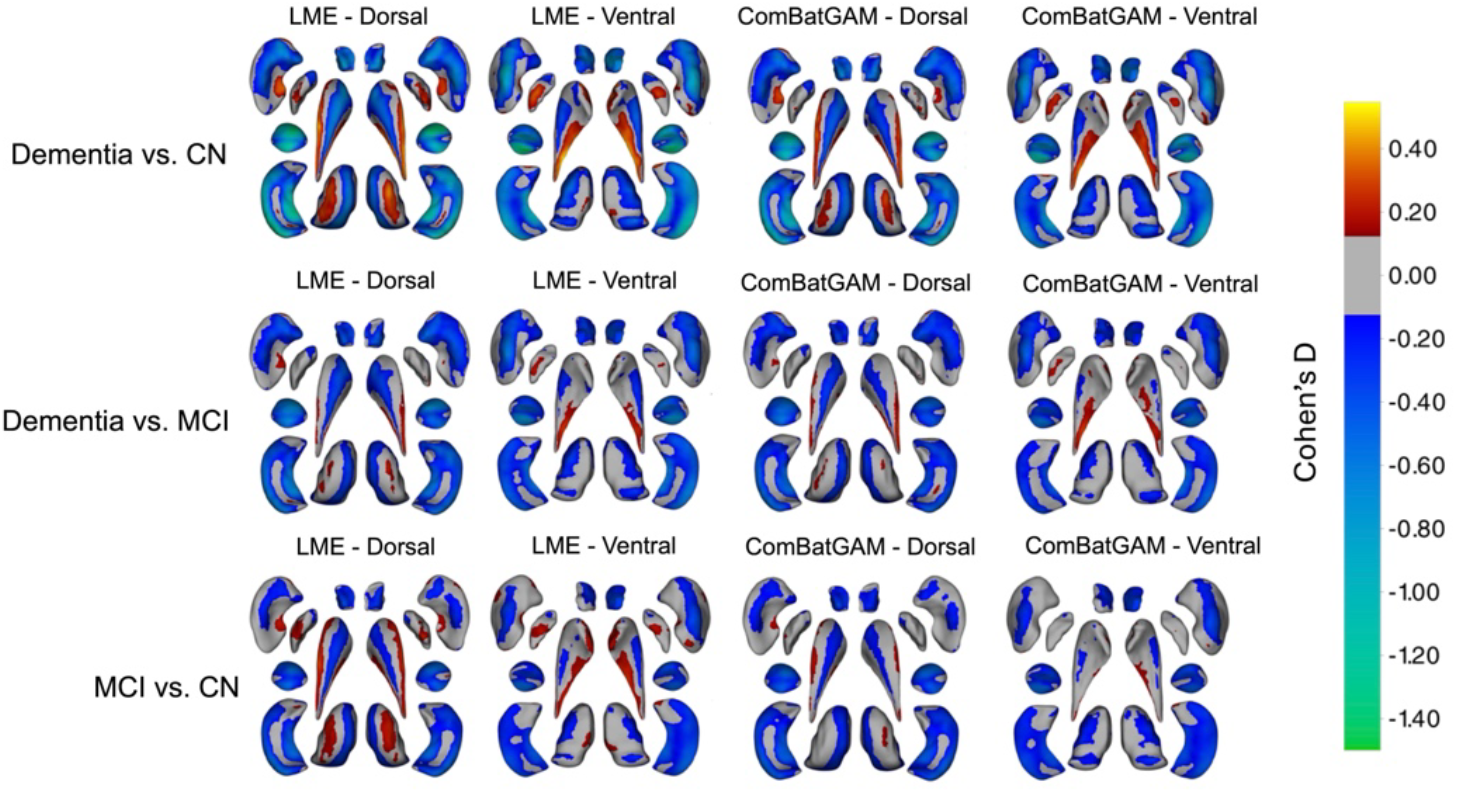
Cohen’s *d* effect size maps for diagnostic group comparisons using both linear mixed-effects models (LME) accounting for site as a random effect, and regression models based on ComBat-GAM adjusted RD shape features. All models were adjusted for age, sex, and intracranial volume (ICV). Warmer (red/yellow) colors indicate greater RD (thickness) and cooler colors (blue/green) indicate lower RD after FDR correction (*q*<0.05) for multiple comparisons. Gray regions indicate vertices not passing statistical significance.

### 3.3 Learning Curve Analysis

Learning curves for each shape-based ML model are shown in **Fig. 4**. For the three models, and across all diagnostic tasks, classification performance increased with training sample size.

**Fig. 4.**
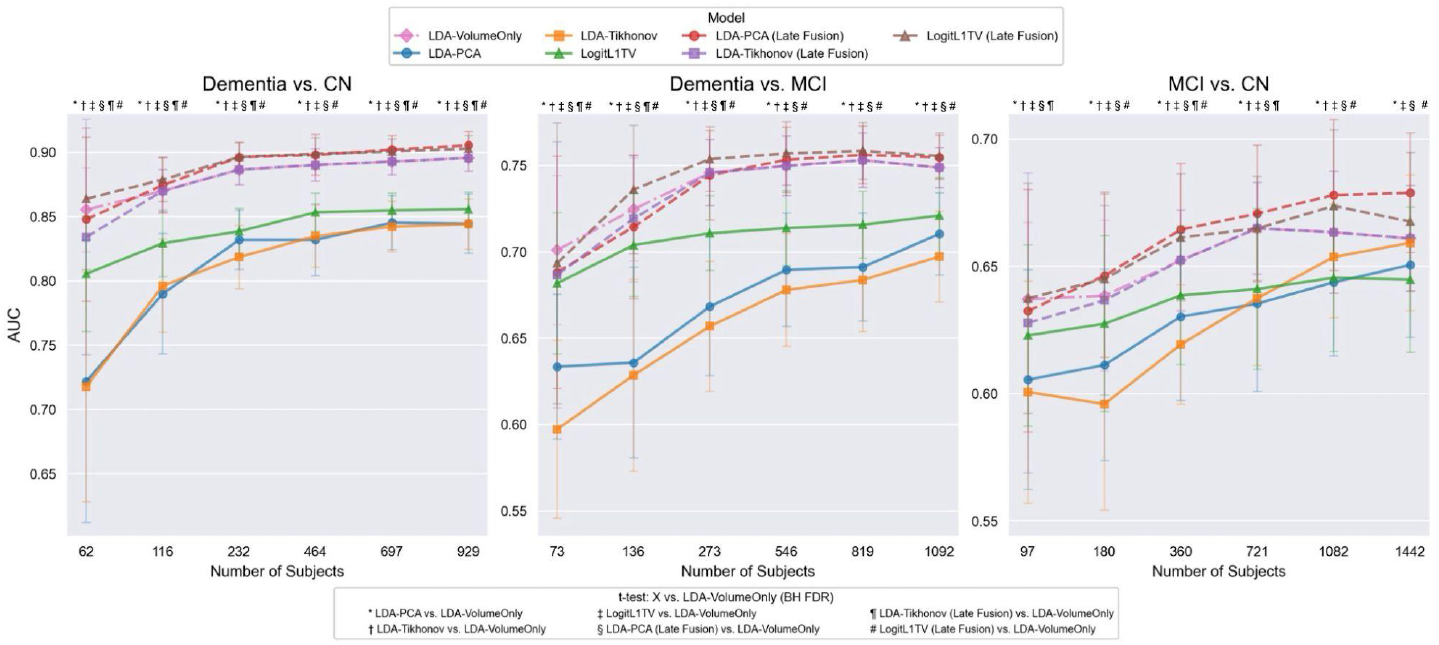
Learning curves are shown (as piecewise linear approximations) for shape-based machine learning (ML) models across three diagnostic classification tasks (dementia vs. CN, dementia vs. MCI, and MCI vs. CN). Average AUC is shown as a function of the number of training subjects where error bars represent standard deviations across the 100 bootstrapped iterations and 5 subsample sizes used for repeated random draws (see detailed methods). Symbols indicate statistically significant differences between shape-based, fusion and volume-based models following FDR corrected t-tests (adjusted *p* < 0.05).

#### Learning Curves Across Diagnostic Tasks

In the dementia vs. CN task, all models achieved relatively high AUC values even at small training sample sizes (AUC=0.72-0.90). Performance increased modestly as additional data were added within this sample range. The dementia vs. MCI classification showed intermediate performance among diagnostic classifications, with AUC values increasing gradually across the evaluated sample sizes (AUC=0.63-0.75). The MCI vs. CN classification was the most challenging task as all models achieved comparatively lower AUC values (AUC=0.60-0.68) compared to other tasks, especially at the smallest training sample size, and performance only increased modestly as additional subjects were included.

#### Comparison of Shape-Based and Late-Fusion Models Relative to the Volume-Based Model

Although the three models exhibited similar learning trajectories, differences in performance were observed for specific diagnostic classifications across certain ranges of training sample sizes (**Fig. 4**). Among the shape-based models, LogitTVL1 achieved the highest numerical AUC, especially for dementia vs. CN and dementia vs. MCI tasks. For MCI vs. CN classification, all three shape-based models showed comparable performance with overlapping confidence intervals. Compared with the shape-based models, the volume-based model generally demonstrated superior classification performance.

Late fusion models numerically outperformed both shape-based and gross volume-based models at all training sample sizes and across all diagnostic tasks. For the dementia vs. CN classification, LogitL1TV-LateFusion achieved the highest AUC at smaller sample sizes, whereas PCA-LDA-LateFusion became the best-performing model at the two largest sample sizes, reaching an AUC of 0.91. For the dementia vs. MCI classification, LogitL1TV-LateFusion ranked first for five of the six evaluated sample fractions, indicating robust performance across varying amounts of training data. For the MCI vs. CN classification, PCA-LDA-LateFusion consistently achieved the highest AUC for all but the smallest sample size, where LogitL1TV-LateFusion showed a marginal advantage.

## 4 Discussion

In this study, we compared the sample-size efficiency of three shape-based, volume-based, and late-fusion ML models for dementia stage diagnosis based on subcortical brain morphometry. As expected, classification performance improved with larger training sample sizes for all models, supporting the general expectation that larger neuroimaging samples improve predictive accuracy. The rate of performance improvement depended more on the task (and underlying diagnostic effect size) than on the model choice. Dementia vs. CN, which showed the largest Cohen’s *d* group differences, consistently achieved the highest AUCs and approached performance saturation with relatively modest sample sizes. In contrast, MCI vs. CN showed slower improvement with increasing sample size, consistent with smaller gross volume and shape differences between groups. These results show that larger underlying feature effect sizes will require fewer training subjects to reach stable performance, while those with smaller effects can continue to benefit from additional training data. This pattern is consistent with scaling-law behavior, in which predictive performance depends jointly on signal strength and sample size rather than model architecture alone.

Among shape models, LogitL1TV generally achieved the highest performance, particularly at smaller training sample sizes. The largest advantage was observed for the dementia vs. MCI classification, where LogitL1TV outperformed LDA-Tikhonov at the two smallest training sample sizes. However, these differences became progressively smaller as additional training data were incorporated, suggesting that the benefit of the TV-L1 regularization is greatest in data-limited settings. Despite these differences, the improvements associated with model selection were smaller than those achieved by incorporating volumetric information through late fusion, showing that feature representation had a greater impact on classification performance than the model selection.

The advantage of LogitTVL1 may reflect differences in how the three models regularize high-dimensional shape features. Tikhonov regularization shrinks model weights independently but does not explicitly model the spatial coherence of the underlying anatomy. In contrast, TV-L1 regularization combines sparsity with spatial smoothness, encouraging neighboring vertices to receive similar model weights while suppressing isolated noisy features. Such a prior is well matched to neurodegenerative conditions such as Alzheimer’s disease, where pathological changes generally extend across contiguous anatomical regions. Despite this advantage, the improvement provided by TV-L1 over the LDA-based approaches was modest. All three methods ultimately learn linear decision boundaries from the same underlying shape measurements, differing only in how those measures are represented and regularized, which may explain why model performance was influenced more by the magnitude of subcortical group differences than by model choice.

The LDA gross volume model outperformed the shape-based models across diagnostic tasks. One possible reason for this is that shape models estimate thousands of correlated vertex-wise features under regularization, which makes them more prone to estimation variance and attenuation of the dominant discriminative signal in limited sample settings. In contrast, gross volume provides a compact, low-variance summary of the dominant atrophy pattern that is more statistically efficient to learn.

Late fusion consistently provided the highest performance across tasks and training sample sizes. This suggests that shape and gross volume features can provide complementary information for improving dementia diagnostic accuracy. Whereas gross volume captures the dominant atrophy pattern, shape features contribute additional complex deformation signals (see **Figure 2** where dementia was associated with thinner and thicker alterations compared to MCI and CN).

Several limitations to the current analysis should be considered: First, this study focused on classification tasks with relatively large and spatially coherent anatomical alterations, which are well studied from the ADNI cohort. Future work should examine conditions with different spatial patterns, including subtler brain effects (psychiatric disorders [9, 10]) or more localized alterations (Huntington’s disease [27]), to determine whether the learning behavior observed in this study generalizes to other illness-related brain signatures. Second, we evaluated only three linear shape-based classification models. Greater differences in scaling behavior may emerge when linear shape-based methods are compared with nonlinear or deep representation-learning approaches, especially across larger ranges of sample sizes. Finally, our analyses were limited to vertex-wise radial distance (local thickness) features from subcortical structures. Other shape features, subcortical surface deformation measures, cortical surface representations, and multimodal imaging metrics (e.g., white matter connectivity, functional MRI) may exhibit different scaling attributes.

In summary, learning curve analysis provides a useful framework for evaluating the scaling behavior of shape-based ML models. By characterizing performance across different sample sizes and classification tasks, learning curves can guide model selection according to both available data and the magnitude of anatomical differences between groups.

## Acknowledgments

This work was supported by NIH grants R21 MH139001, R01 MH129742, R01 MH131806, R01 AG058854, R01 MH134962. Research reported in this publication was supported by the Office of The Director, National Institutes of Health under Award Number S10OD032285. The content is solely the responsibility of the authors and does not necessarily represent the official views of the National Institutes of Health. Data collection and sharing for this project was funded by the Alzheimer’s Disease Neuroimaging Initiative (ADNI) (National Institutes of Health Grant U01 AG024904) and DOD ADNI (Department of Defense award number W81XWH-12-2-0012).

## Requirements and Code Availability

The scripts used for this paper will be included in the Brain Geometry Toolkit (v1.1) which is compatible with Python 3.8.x in major operating systems such as Mac OS X, Windows, and Linux. Our code requires independent installation of the following Python packages: pandas (v2.0.x), numpy (v1.24.x), and scikit-learn (v1.3.x). Development is ongoing to incorporate additional modules that accommodate additional types of shape data and AI models. All code is available upon request. Once it is fully validated, it will be made publicly accessible on the ENIGMA GitHub repository (https://github.com/ENIGMA-git).

